# Imperative role of particulate matter in innate immunity during RNA virus infection

**DOI:** 10.1101/2020.03.28.013169

**Authors:** Richa Mishra, K Pandikannan, S Gangamma, Ashwin Ashok Raut, Himanshu Kumar

## Abstract

Sensing of pathogens by specialized receptors is the hallmark of the innate immune response. Innate immune response also mounts a defense response against various allergens and pollutants including particulate matter present in the atmosphere. Air pollution has been included as the top threat to global health declared by WHO which aims to cover more than three billion people against health emergencies from 2019-2023. Particulate matter (PM), one of the major components of air pollution, is a significant risk factor for many human diseases and its adverse effects include morbidity and premature deaths throughout the world. Several clinical and epidemiological studies have identified a key link between the PM composition and the prevalence of respiratory and inflammatory disorders. However, the underlying molecular mechanism is not well understood. Here, we investigated the influence of air pollutant, PM_10_ during RNA virus infections using highly pathogenic avian influenza (HPAI). We thus characterized the transcriptomic profile of lung epithelial cell line, A549 treated with PM_10_ prior to infection with (HPAI) H5N1 influenza virus, which is known to severely affect the lung and cause respiratory damage. We found that PM_10_ regulates virus infectivity and enhances overall pathogenic burden in the lung cells. Additionally, the transcriptomic profile highlights the connection of host factors related to various metabolic pathways and immune responses which were dysregulated during virus infection. Overall our findings suggest a strong link between the prevalence of respiratory illness and the air quality.

## INTRODUCTION

Seven million people are estimated to be killed every year by the air pollution according to the WHO (http://www.who.int/mediacentre/news/releases/2014/air-pollution/en/). WHO has recommended standard permissible level of air contaminants but nearly 80% of the urban cities are well above the standard permissible level (https://www.who.int/airpollution/data/cities/en/). Alarming rate of air pollution in recent years is known to be linked with increased mortality rate and affected the global health and economy [1-6]. One of the major components of air pollution is particulate matter (PM). PM collected from different sources or geographical area may have different impact on the inflammatory and innate immune responses corresponding to the virus infection on human health. Airborne PM were considered the hazardous causative determinants of several diseases such as respiratory, cardiovascular and neurological disorders. These particles are divided into three main categories on the basis of their diameter: coarse particles, or PM_10_, (with an aerodynamic diameter between 10 and 2.5 µm); fine particles, or PM_2.5_, (with diameters < 2.5 µm); and ultrafine particles, or PM_0.1,_ (with diameters < 0.1 µm) [7]. Numerous studies revealed that particulate matter collected from different locations all over the world is strongly associated with the elevated morbidity and mortality and various diseases [8-13]. Several studies have attempted to understand the link between PM isolated from heavily populated regions of India and associated health concerns in term of occurrence of disease [14-19]. Although, most of the studies were based on the epidemiological data and cross-sectional studies, there were few studies about involvement of PM in respiratory diseases [20-22], asthma [23], cancer [24-27], tuberculosis [28, 29]. It has been known that PM can induce innate immunity and can change the level of cytokines, upon its exposure to the airways of humans [30-33]. PM were readily associated with respiratory infections such as chronic obstructive pulmonary disease (COPD) [34-37] and it is also reported to be associated with the respiratory syncytial virus (RSV) and influenza virus infections. [38-41]. Yet these studies are limited to epidemiological, cross-sectional studies [22, 42-44].

Here, we isolated and characterized PM_10_ from a heavily industrialized city Bengaluru, India and checked its effect on RNA virus infection. We observed and concluded that PM_10_ hijacks the innate immune system upon viral infection and significantly enhanced the viral replication of the RNA viruses like new-castle disease virus (NDV), influenza virus - H1N1 (PR8) and H5N1. By performing RNA sequencing analysis, we found that pre-exposure of PM_10_ to the cells downregulates the anti-viral innate immunity related genes in lung (A549) cells during H5N1 infection. Additionally, we reported the upregulation of some previously unknown metabolism-related genes by global transcriptomic profile analysis and observed its role during virus infection as demonstrated by knock down studies of identified genes. These metabolic-related genes play significant role in promoting viral replication in presence of airborne PM.

## RESULTS

### Physical and chemical characterization of PM_10_

To investigate the airborne particles, precisely known as coarse size particulate matter (PM_10_), that were collected and used in the study, we performed SEM-EDS analysis of PM_10_ collected from Bengaluru city, India. SEM-EDS techniques decipher the particle shape and chemical composition. It is a method for high resolution surface imaging using electron beams. SEM-EDS analysis provided us an understanding about the differences in morphology and elemental composition of the airborne PM_10_ collected samples. To understand the effect of PM_10_ on host cells, we initially characterized the particles through imaging and identified that various shapes were embedded in the particulate matter. We found different biologically active morphological features within the particulate matter PM_10_ (Fig. 1). These varied characteristic features of PM_10_ consists of biologically active shapes like air ash, spherical, irregular, well-defined, aggregates and rounded. Next, we investigated the types and concentration of elements present in PM_10_ to decipher the origin in terms of biogenic, geogenic and anthropogenic particles. To this end, we performed energy dispersive spectroscopy (EDS) analysis and found different concentrations of various metals. We got different peaks in the spectrum obtained upon analysing the sample at different points with the pulse of electrons (Supplementary Fig. S1A). The peaks in the spectra correspond to the presence of different elements particularly metals (% by weight) in the particulate matter (Supplementary Fig. S1B). Some of the listed metals and non-metals (in traces and/or abundance) are iron, carbon, oxygen, aluminium, lead, silver, silica, titanium, cadmium, sodium, chloride, magnesium, copper, zinc, gold, tin, vanadium, chromium, nickel, arsenic, molybdenum, barium, potassium, sulphur, strontium, manganese, cobalt and selenium.

**Figure 1:**
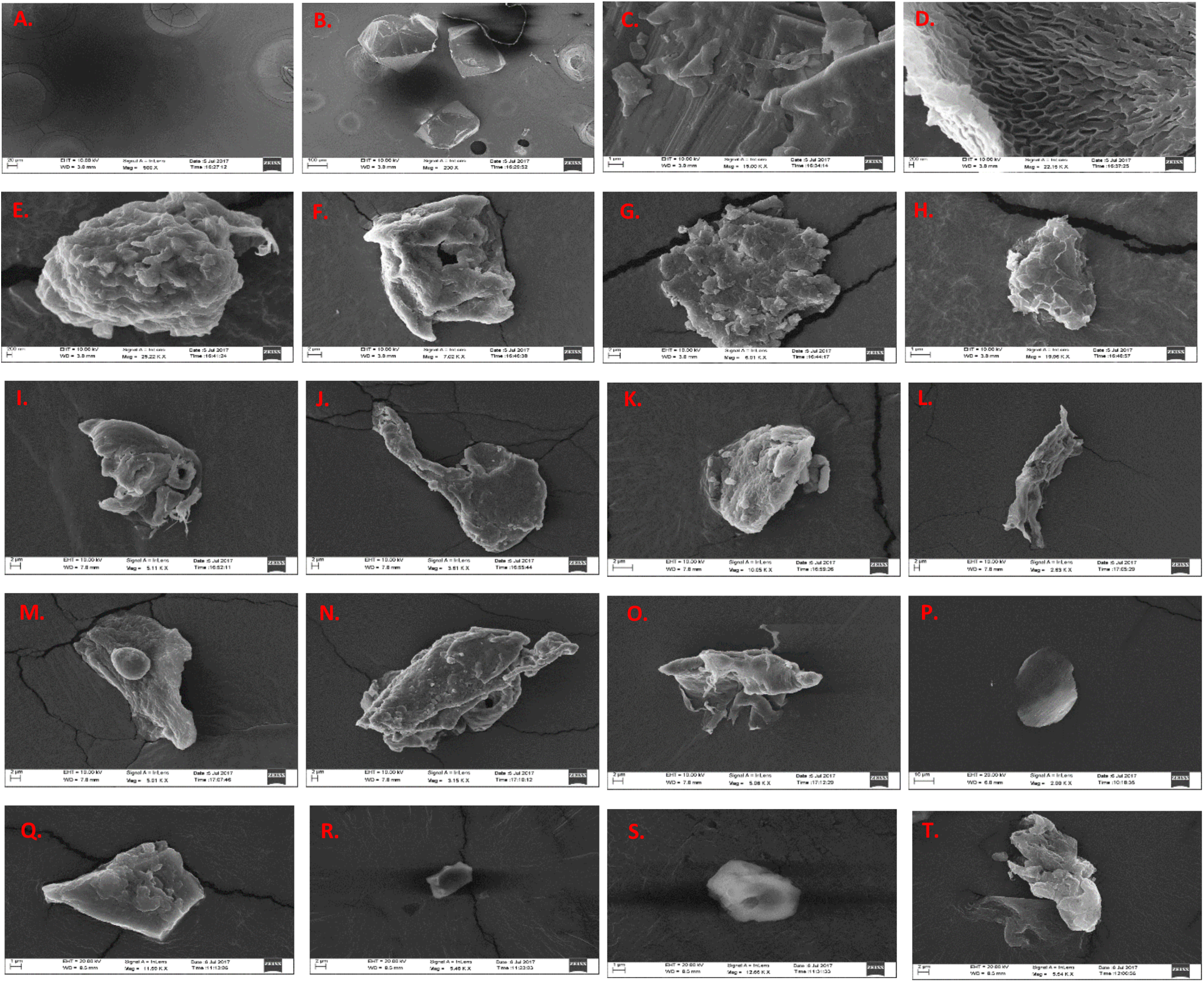
Morphological features of PM_10_. Scanning electron images of coarse airborne particulate matter PM_10_. Image of blank solution with alone with no PM dissolved in it. (B-T) Images of different shapes with varied structures representing the different characteristic morphological features of PM in the samples.

### Exposure of PM_10_ reduces innate immunity upon RNA virus infection

As reported previously, particulate matter or similar substances like smog, diesel exhaust, cigarette smoke extract causes activation of the inflammatory responses when comes in contact with host’s airways and lungs [41]. Therefore, characterization of PM_10_ prompted us to examine whether PM_10_ can induce any innate immune responses in human lung epithelial carcinoma cells, A549. Interestingly, we have found that when cells were exposed to PM (II), which correspond equal volume of PM_10_ and DMEM media, type I interferon, IFNβ (Fig. 2A) and inflammatory cytokine IL-6 (Fig. 2B) were induced. Furthermore, we performed IFNβ and ISRE promoter assay after infection with NDV in presence of PM_10_ and found that there was significant reduction in the promoter activities at the dosage of PM_10_ (II) (Fig. 2C). Additionally, we concluded that in different set of experiments dual treatment of PM_10_ and virus infection (NDV) to A549 cells as shown in schematic representation (Fig. 2D) reduces the mRNA transcript levels of interferon IFNβ and cytokine IL-6 (Fig. 2E-F). These findings further prompted us to investigate whether currently characterized PM_10_ is associated with any respiratory diseases because majority of infectious respiratory diseases are mainly caused by RNA viruses.

**Figure 2:**
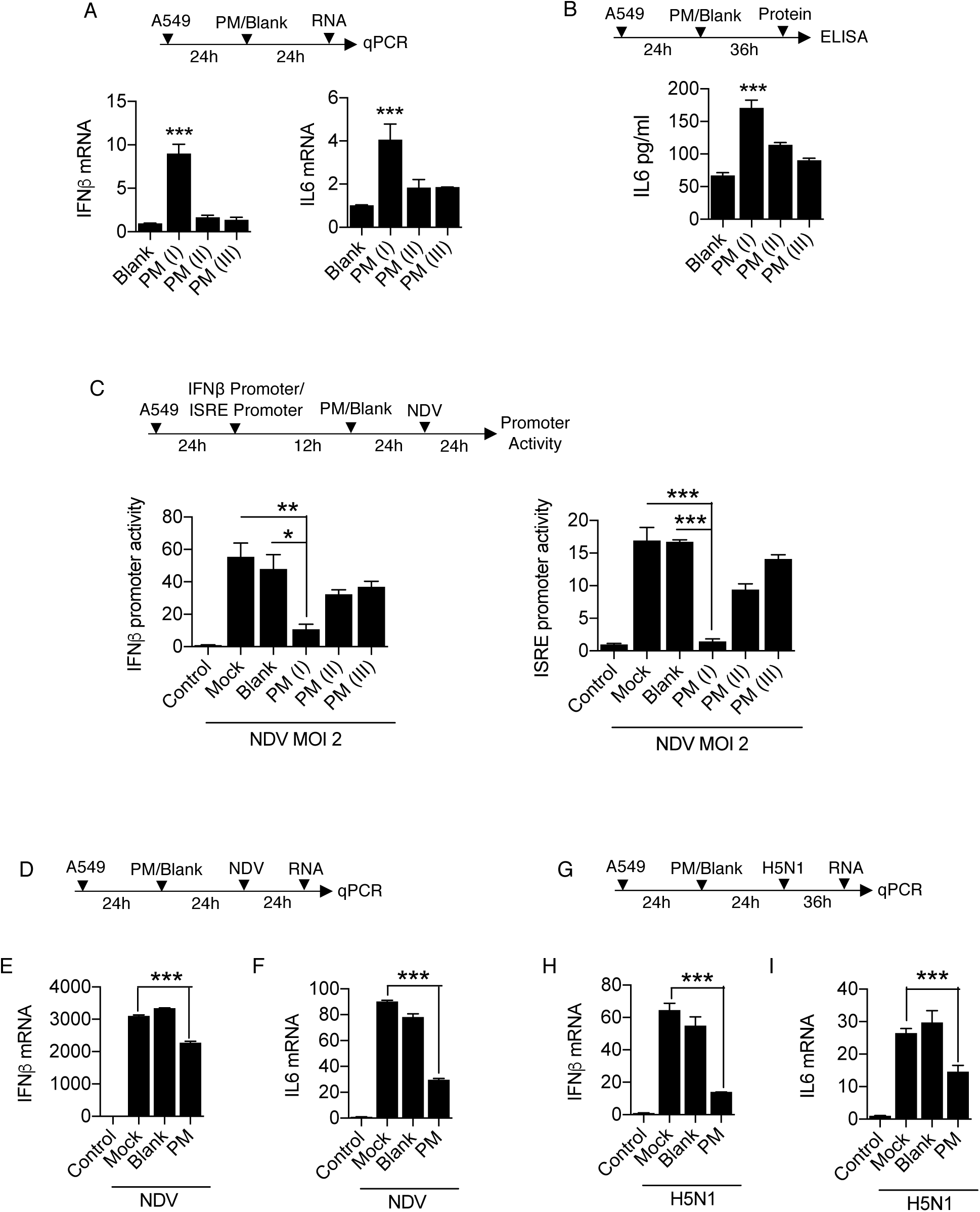
PM_10_ regulates the innate immune response upon RNA virus infection. Quantification of innate immune response. A549 cells were treated with PM_10_ and control mentioned as blank for (A) 24 hours then harvested in Trizol to quantify the mRNA expression of *IFNβ* and *IL6* by qRT-PCR. (B) 36 hours then cell supernatant was collected to measure the protein level of *IL6* by ELISA. (C) Schematic representation of workflow for quantification of IFNβ and ISRE promoter activities by luciferase assay as indicated in A549 cells. NDV represents new-castle disease virus infection at MOI = 2. (D) Schematic work flow of PM_10_ exposure and NDV infection. (E) Quantification of *IFNβ* and *IL6* mRNA transcripts in uninfected (control), mock infected, blank treated and PM_10_ exposed cells by qRT-PCR. (G) Schematic work flow of PM_10_ exposure and H5N1 influenza infection. (H) Quantification of *IFNβ* and *IL6* mRNA transcripts in uninfected (control), mock infected, blank treated and PM_10_ exposed cells by qRT-PCR. Data are mean +/- SEM of triplicate samples from single experiment and are representative of two independent experiments. \*\*\**P*<0.001, \*\**P*<0.01 and \**P*<0.05 by one-way ANOVA Tukey test and unpaired t-test.

Previously, it has been shown that cigarette smoke extract (CSE) affects various regulatory pathways during rhinovirus (RV) infection using human bronchial cell lines by microarray analysis [41]. We re-analyse the GEO dataset: GSE27973 in context to our prospective and found that there are several important cellular machineries associated genes (Supplementary Fig. S2A) were modulated due to CSE exposure and RV infection. We next analysed the regulation of important genes involved in diseases particularly influenza (flu) virus infection and key immune signalling pathways (Supplementary Fig. S2B-D). Gene profile analysis concluded that various antiviral genes were prominently downregulated upon CSE exposure and RV infection. Here, in current study we used influenza virus infection along with PM_10_ treatment in the A549 cells, because influenza virus infection is severely fatal compared to any other virus that causes respiratory damage and influenza virus is regularly active upon the evolutionary scale and regarded as one of the hazardous threats according to WHO to humans. Therefore, to get insights about PM_10_ exposure and highly pathogenic avian influenza infection (HPAI), we treated the A549 cells with PM_10_ and infected them HPAI H5N1 (MOI 2) as shown in schematic representation (Fig. 2G). We observed that that PM_10_ reduces the mRNA expression levels of both IFNβ and IL-6 in presence of H5N1 infection (Fig. 2H-I), indicating that during pathogenic infection by RNA viruses, particularly influenza virus, PM_10_ reduces the innate immune response in the cells.

### PM_10_ enhances viral replication upon RNA virus infection

Curtailed immune responses upon PM_10_ treatment and virus infections: both in case of NDV and H5N1 influenza virus infections, prompted us to measure the viral load in presence of PM_10_. We thus demonstrated the experiment of PM_10_ exposure and virus infection like NDV, H1N1 (PR-8) and H5N1 in A459 cells respectively. Using virus-specific primer, it was observed that PM_10_ significantly enhances the viral replication of all the RNA viruses ubiquitously. PM_10_ enhances the virus replication of NDV (Fig. 3A), H5N1 (Fig. 3B) and H1N1 (Fig. 3C). Additionally, microscopy analysis demonstrates similar results in which GFP tagged NDV was used to infect the PM_10_ pre-exposed cells (Fig. 3D). Increased NDV infection was quantified by measuring the intensity of GFP signal and number of GFP positive cells (Fig. 3E-F). Furthermore, presence of PM_10_ along with NDV infection induces cell death as an additional detrimental effect on cells, quantified by the trypan blue assay (Fig. 3G). Altogether, our results conclude that PM_10_ enhances the viral replication pertaining to lower immune responses.

**Figure 3:**
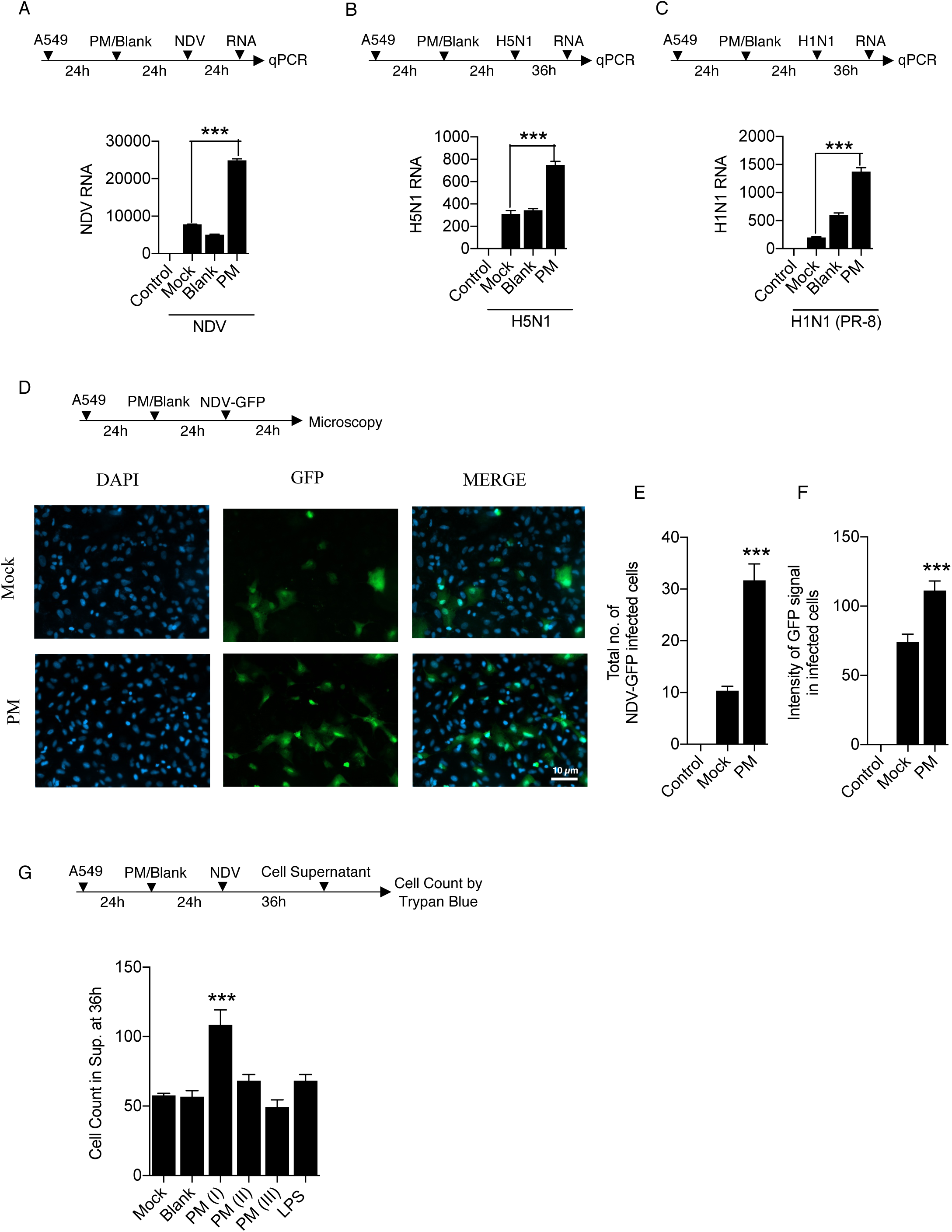
PM_10_ elevates the RNA virus infection. (A-F) Estimation of viral replication in A549 cells exposed with PM_10_ for 24 hours before virus infection at MOI = 2. (A) Schematic work flow of the experiment, PM_10_ enhances the NDV abundance (viral transcripts) in the cells compared to control groups (uninfected control, mock infected, blank treated and PM_10_ exposed). (B-C) Schematic work flow of the experiment, PM_10_ enhances the H5N1 and H1N1 abundance (viral transcripts) in the cells compared to control groups (uninfected control, mock infected, blank treated and PM_10_ exposed). (D) Schematic work flow for microscopy: A549 cells were exposed with PM_10_ then after infected with GFP – labelled NDV for 24 hours, cells in the cover slips were then fixed as per the protocol described in methods section and estimated for GFP positive signals quantified as (E) total number of NDV-GFP infected cells and (F) intensity of GFP signals in infected cells. (G) Schematic work flow to estimate the cell death in cell supernatant after PM_10_ exposure and NDV infection in A549 cells. Cells (dead) were counted by the trypan blue counting assay. Data are mean +/- SEM of triplicate samples from single experiment and are representative of two independent experiments. \*\*\**P*<0.001 by one-way ANOVA Tukey test and unpaired t-test.

### RNA-Seq analysis of H5N1 infected cells in presence of PM_10_

PM_10_ enhances the viral replication and suppress the immune responses. To further understand the global outcome of immune responses within the human cell and to dissect the mechanism about the current physiological effect, we performed RNA sequencing to profile the overall changes in the host genes and cellular pathways upon PM_10_ treatment and HPAI H5N1 infection. Schematic workflow of the experiment and transcriptomic sequencing shown in Fig. 4A. Differential expression of host genes analysis was performed between PM_10_-treated_H5N1-infected and subsequently mock-treated_H5N1-infected samples. Differentially expressed genes were marked in red and other regulated genes which were altered more than 1.5 fold were marked in blue, altogether they were represented by a volcano plot (Fig. 4B). Next, to understand the overall cellular changes, gene ontology analysis was performed through DAVID tool to obtained the enriched biological terms from the top differentially expressed genes with the fold change between -1.5< log FC >1.5. The top enriched pathways were depicted in bubble plot and circle plot generated through R package GOplot (Fig. 4C). Herewith, bubble plot represents the significant enriched ontology terms like biological process (BP), cellular components (CC) and molecular functions (MF). Circle plot represents the connection between these significantly enriched ontology terms and the status of genes contributing to each ontology terms. Additionally, the chord plot represents the connection of common significant differentially expressed genes with the significant enriched ontology terms (Supplementary Fig. S3A). Gene ontology analysis revealed that significantly down-regulated genes during H5N1 infection in presence of PM_10_ were involved majorly in various immune signaling pathways and innate immune responses, in accordance with our experimentally validated results. On contrary, comprehensive analysis revealed that significantly up-regulated genes were majorly involved in various metabolic pathways. To test this, pathway enrichment analysis was performed through DAVID tool and top enriched pathways of differentially expressed genes with -1.5<log FC<1.5 were represented by the chord plot depicting the network between significant differentially expressed genes and their enriched pathways. Additionally, circle plot depicts the connection of top enriched pathways with the status of the genes contributing to the pathway represented by their logFC and Z-score (Fig. 4D). Furthermore, representative of up-regulated genes from significantly regulated metabolic pathways were validated by qRT-PCR analysis and found the enhanced mRNA expression levels of VIPR1, CYP1A1, AlDH1A3 and PPP1R14A genes upon H5N1-infection in A549 cells in presence of PM_10_ (Fig. 4E). Similar results were obtained in NDV-infected A549 cells in presence of PM_10_ (Supplementary Fig. S3B-E). Related results were obtained by re-analysing the GEO dataset GSE27973 of rhinovirus infection and CSE exposure in human bronchial epithelial cell lines (Supplementary Fig. S4A-B). Additionally, theses metabolic pathways-related genes were found to be associated with many pathological states (Supplementary Fig. S4C). Overall our data concludes that upon PM_10_ treatment during RNA virus infection, particularly, influenza virus infection, PM_10_ significantly enhances the virus infection by down-regulating innate immune responses and upregulating different metabolic processes, that might cater air pollutant to enhance virus infectivity within the cells and manifold enhance respiratory damage.

**Figure 4:**
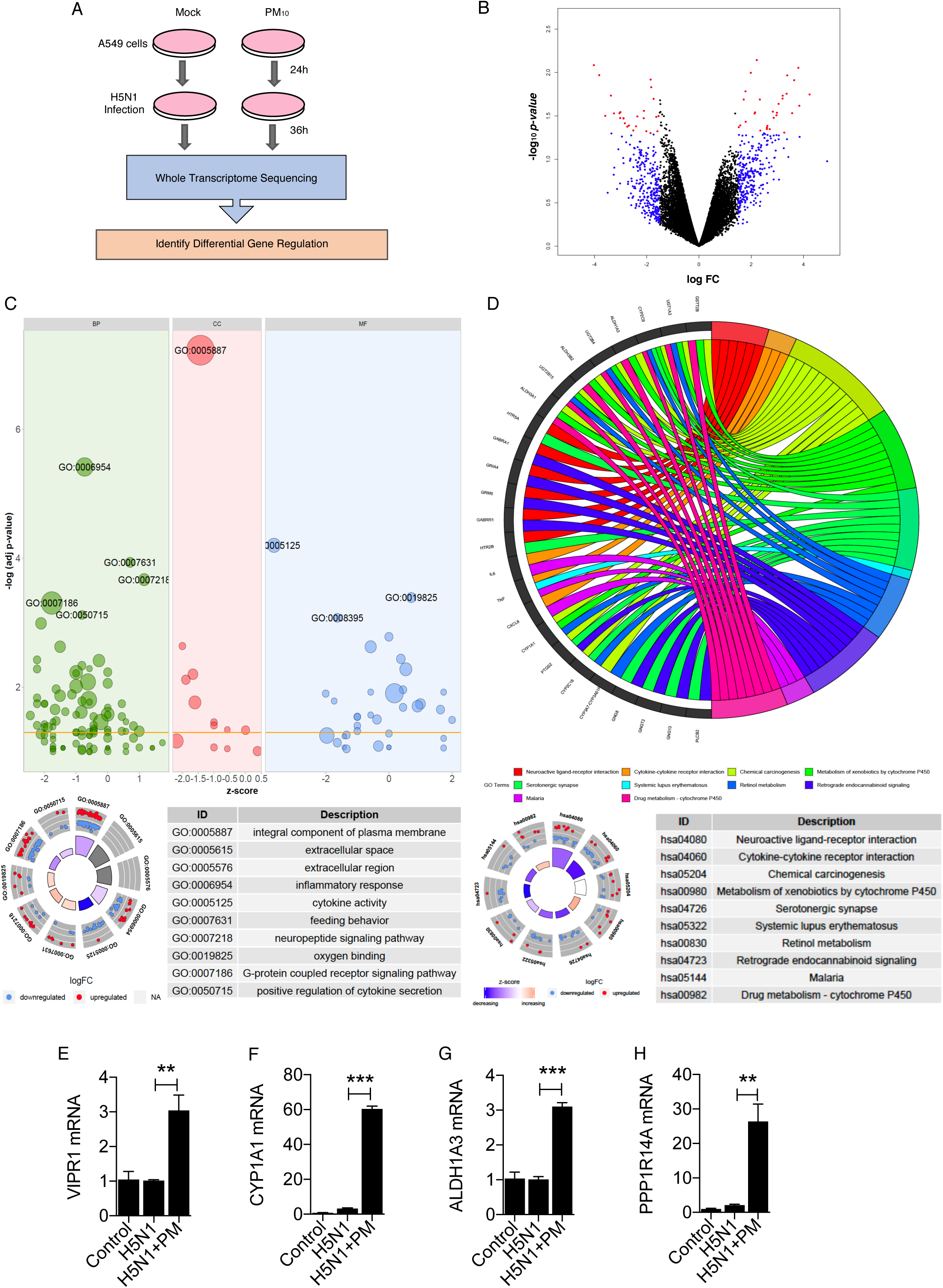
Transcriptomic analysis shows PM_10_ enhances abundance of metabolic pathways-related transcripts (genes) during H5N1 infection. (A) Schematic outline of PM_10_ exposure and H5N1 infection (MOI 2) in A549 cells at indicated time. Cells were subjected to whole transcriptome sequencing and differential gene expression analysis. Volcano plot represents differential expression of genes between two groups of samples (mock H5N1 infected and PM_10_ exposed plus H5N1 infected) during H5N1 infection in A549 cells. For each gene: *P-value* is plotted against fold change (mock vs PM_10_). Significantly differentially expressed genes are marked in red colour while genes which are altered (>1.5-fold) are marked in blue colour. (C) Gene Ontology analysis performed as per the protocol mentioned in methods section represents the top differentially expressed genes in ontology terms: BP (biological processes), CC (cellular components) and MF (molecular functions) respectively depicted by bubble plot and circle plot generated through R package GOplot. (D) Pathway enrichment analysis performed as per the protocol mentioned in methods section. Chord plot represents the differentially expressed genes and their connection with the top enriched pathways. Circle plot represents the top enriched pathways and status of the genes contributing to the pathways by their logFC and Z-score. (E-H) Quantification (measured by qRT-PCR) and validation of the fold changes in the abundances of significantly expressed metabolic pathways related transcripts: VIPR1, CYP1A1, ALDH1A3 and PPP1R14A in the samples of A549 cells; untreated (control), mock H5N1 infected (H5N1) and PM_10_ exposed plus H5N1 infected (H5N1+PM), analyzed by RNA-Sequencing. For figure (E-H): Data are mean +/- SEM of triplicate samples from single experiment and are representative of two independent experiments. \*\*\**P*<0.001 and \*\**P*<0.01 by one-way ANOVA Tukey test and unpaired t-test.

### Knockdown of metabolism-associated genes involved in virus replication

To investigate the correlation between the upregulated metabolic pathways-related genes and their influence on virus infection upon PM_10_ treatment, we selected CYP1A1, VIPR1 and PPP1R14A genes because these genes were significantly upregulated in our RNA sequencing analysis and were their role is poorly understood. The CYP1A1 involved in xenobiotic metabolic pathways, which is one of the metabolic pathways aiding virus infections, VIPR1 is associated with G-protein coupled receptor pathway and PPP1R14A involved in vascular smooth muscle contraction and oxytocin pathway which were directly or indirectly related to virus infectivity within the host cell. To this end, we performed knockdown study of CYP1A1, VIPR1 and PPP1R14A in A549 cells. We used two different short hairpin (*sh)*-clones for each gene to knockdown the expression of CYP1A1, VIPR1 and PPP1R14A genes respectively as shown in the schematic workflow (Fig. 5A-C). Particularly, knock down of these genes in presence of NDV infection in A549 cells, leads to significant suppression the virus infection, notably, the knockdown substantially reduced the gene expression (Fig. 5A-C) suggesting that upregulated metabolic pathways-related genes in presence of airborne particulate matter (PM_10_) support virus infections that further contribute to the severity of respiratory related diseases or highly pathogenic respiratory virus infections, like influenza.

**Figure 5:**
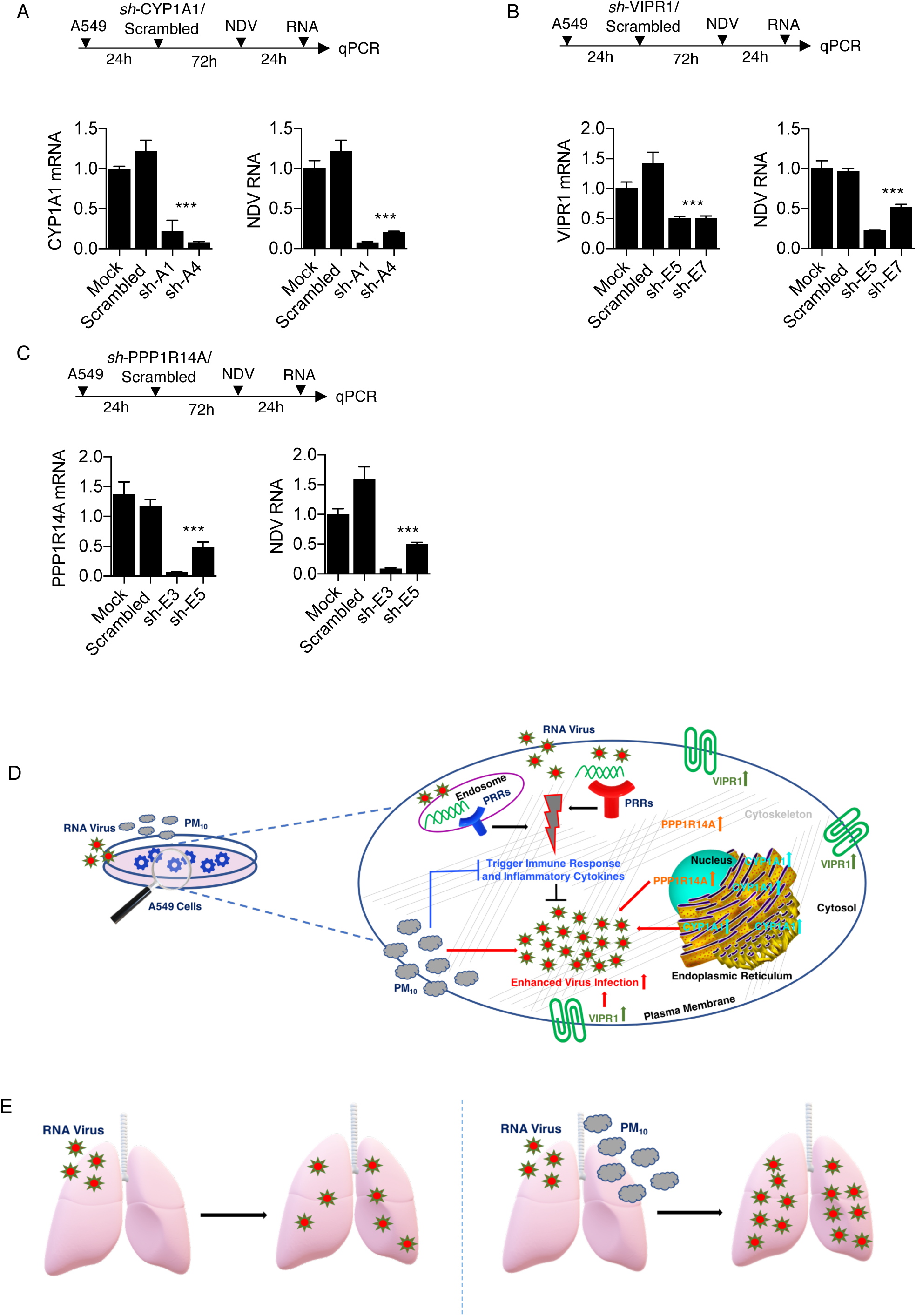
Knockdown of validated genes reduces RNA virus infection. A549 cells were transiently transfected with 1.5μg of two respective *sh-*clones of each indicated genes or scrambled control for 72 hours then infected with NDV (MOI 2) for 24 hours and subjected to the quantification of the NDV viral RNA transcripts and the respective indicated transcripts or genes (A) CYP1A1, (B) VIPR1 and (C) PPP1R14A. (D) Graphical representation of the study: CYP1A1, PPP1R14A and VIPR1 at their respective location (endoplasmic reticulum, cytoskeleton-nucleus and plasma membrane respectively) within the cell induced upon RNA virus infection and PM_10_ exposure to increase viral infection in presence of airborne PM_10_. PRRs – Pattern Recognition Receptors to sense the viral particles. Overall immune responses were downregulated in PM_10_ treated cells. (E) Cumulative effect of PM_10_ and virus infection enhance respiratory damage and overall virus infection in lungs at the organismic level.

## DISCUSSION

In modern world, air pollution and emergence of novel microbial pathogens infecting through respiratory route has been included as the top threat to global health in the 13th General Programme of work, by WHO which aims to cover more than three billion people against the health emergencies from 2019-2023. (https://www.who.int/about/what-we-do/thirteenth-general-programme-of-work-2019-2023). Air pollution is a one of key risk factor for respiratory route or metabolism-associated diseases, and its adverse effects include morbidity and premature deaths throughout the world [45]. Particulate matter contributes to the majority of lethal effects caused by air pollution, which differs according to the geographical area. Particularly in India, where air pollution is predominant factor in major cities like, New Delhi, Bengaluru, Pune and so on. There were so far, very fewer studies which links particulate matter with the health and immunity in context to respiratory virus infections [37, 46]. Air pollutants are one of the major health concerns especially in inducing the adverse effects during pathogenic infections. Though these pollutants modulate the host defense and enhance susceptibility and severity during infection, the underline mechanisms are poorly understood [47]. Influenza is also included among the topmost threats by WHO and suggested to have pandemic potential. Influenza infection peaks during the winter season and cause frequent seasonal endemics, as well as sudden unforeseen pandemics. It spreads readily, and there is no proper vaccination available, therefore, it’s been a major health as well as an economic burden throughout the world. The factors contributing to the emergence of a sudden pandemic strain of influenza is not well understood. Environmental factors play an essential role in the severity and spread of respiratory infections particularly influenza infection. Few studies explained the direct causative effects of ambient pollutants and other similar causative agents like cigarette smoke extracts, diesel exhaust on various lung infections and especially on the severity of common cold occur by rhinovirus [37, 41, 48]. Different studies provide varied results over the impact of particulate matter in lung infections, as they are from different geographical origins [44, 49, 50]. In a developing country like India, the level of ambient airborne particulate matter, especially PM_10_, increased in the past decade due to heavy industrialization. PM_10_ isolation from Indian subcontinent and its deleterious effects on human health in context to hampering the innate immune defense, against RNA virus infections are not reported yet.

Herewith in this particular study, we sought to understand whether PM_10_ exposure leads to significant modification of innate immune responses and viral infectivity in human lung epithelial cell lines, A549. Additionally, we focused to explore the overall cellular changes occur when cells were exposed to PM_10_ and virus infection together. We also aimed to underpin the mechanism behind the intensification of influenza (H5N1) virus and other RNA virus infections like NDV in presence of airborne particulate matter (PM_10_). These cellular outcomes persuaded us to perform the RNA sequencing and analyse transcriptomic profile to unravelled the cellular changes during PM_10_ exposure during infection.

We used PM_10_ in our study obtained from Bengaluru city. Bengaluru is one of the heavily industrialized area in India. Therefore, studying the characteristics of ambient particulate matter around Bengaluru area is of importance. Initially, we characterized the PM_10_ by performing SEM-EDS analysis, and reported the morphological features and chemical composition of the particulate matter as revealed by imaging analysis. PM_10_ and its impact on airway was investigated by exposing the cells with PM_10_ and infecting them with different RNA viruses like NDV and H5N1 flu virus. Our results demonstrate the consequences of both air pollutant and virus infection. Interestingly, we observed that PM_10_ isolated from the Bengaluru demonstrate that PM_10_ suppresses innate immunity and significantly elevate viral replication. Previously, it has been shown that antiviral response was supressed upon CSE exposure during rhinovirus infection in human bronchial epithelial cell lines [41]. This prompted us to test the effect of PM_10_ on the enhanced infectivity of highly pathogenic avian H5N1 influenza infection and decipher the molecular mechanism.

Although, few studies are reported the global transcriptomic changes, in presence PM_10_ by microarray analysis. We, for the first time, used high throughput RNA sequencing to study the overall changes in the gene expression upon PM_10_ exposure during the viral infection of highly pathogenic avian Influenza (HPAI) H5N1 virus in the lung carcinoma cells, A549. RNA sequencing analysis identified that majority of genes are significantly downregulated were involved in immune-related pathways, cytokine signalling, and few other inflammatory pathways. In addition to this, we observed a significant increase in the expression of genes involved in various metabolic pathways, which were previously remain unknown, particularly in air pollution. We validated RNA sequencing results for four of the top hits genes namely VIPR1 (vasoactive intestinal peptide 1), CYP1A1 (cytochrome P450, family 1, subfamily A memeber1 also known as aryl hydrocarbon hydroxylase), ALDH1A3 (aldehyde dehydrogenase 1, family member 3A) and PPP1R14A (protein phosphatase 1 regulatory inhibitor subunit 14A) using quantitative qRT-PCR analysis. These selected genes are, VIPR1, mainly located on plasma membrane and PPP1R14A majorly located on nucleus and cytoskeleton were moderately found to be involved in virus infections like HIV-1 and influenza as reported by an *in-vitro* study and an *in-silico* phosphoproteomics study in human macrophages respectively [51-53]. CYP1A1 was recently reported to be involved in many virus infections especially hepatitis B and hepatitis C virus [54-56]. One such report superficially uncovers the induction of CYP1A1 in presences of PM_10_ [57]. Additionally, induction of CYP1A1 in presence of diesel exhaust particles were extensively reported in human bronchial cells [58]. Apart from studies related to different types of cancers [59, 60], ALDH1A3 was also previously reported in connection with virus infections like human papilloma virus and respiratory syncytial virus [61-63]. Altogether, these significant differentially expressed genes noted in our study related to different metabolic modifications inside the cell and reasonably linked to virus infections, therefore, we selected these genes for validation in context to RNA virus infectivity. We demonstrated by *sh*-RNA mediated transient silencing that these genes significantly reduced the viral replication. This states the importance of these metabolic pathway-related genes in regulation of pathogenic burden during viral infection.

Overall, this study highlights the effect of PM_10_ exposure upon virus infection that affects the lung airways to cause severe respiratory damage. And high throughput RNA sequencing was performed for the first time, in context to Indian subcontinent distribution of particulate matter (PM_10_). PM_10_ collected and isolated to study the transcriptomic changes upon its exposure during influenza infection in A549 cell lines. The overall summary of the study was graphically illustrated in Figure 5D-E. There were very few studies that reported the link between PM_10_ exposure and enhanced viral infections [64, 65]. Our study not only reported the status of viral replication upon PM_10_ exposure, but also examined the role of metabolic pathways - associated genes involved in the viral replication. Still, this study requires further *in-vivo* analysis using mice models in order to explore the effect of pollutant under physiological condition after PM_10_ exposure. Further studies were needed to uncover the connecting links between other respiratory infectious diseases and the use of PM_10_ from different geographical locations, seasonal variation, which will give better insights about the effects of PM_10_ over various lung infections including influenza virus infection.

## MATERIALS AND METHODS

### Cell lines and reagents

A549 human alveolar basal epithelial cells (Cell Repository, NCCS, India) were cultured in Dulbecco’s modified Eagle’s medium (DMEM) supplemented with 10% fetal bovine serum (FBS) and 1% Antibiotic-Antimycotic solution. DMEM, FBS and Antibiotic-Antimycotic solution were purchased from Invitrogen. Ambient particulate matter of coarse particle size PM_10_ was obtained from Dr. Gangamma S. which was collected and isolated in appropriate solvent media from the geographical regions of Bengaluru city, at NITK, Surathkal, Mangaluru, Karnataka. A549 cells were seeded in 12 well culture plate at a concentration of 3×10^5^/well overnight (37°C, 5% CO_2_). Cells were treated with PM_10_ along with controls namely blank and/or LPS (100 ng) for 24 hours prior to infection. Plasmids containing Firefly Luciferase gene under *IFNβ* and *ISRE* promoters, were obtained from Professor Shizuo Akira’s (Osaka University, Japan). All sh-clones, were obtained from the whole RNAi human library for shRNA mediating silencing (Sigma, Aldrich) maintained at IISER, Bhopal, India.

### Virus Infection

Airborne particulate matter (PM_10_) treated A549 cells were infected with new-castle disease virus (NDV), highly pathogenic avian influenza virus (H5N1) and vaccine strain PR-8 virus (H1N1) at respective multiplicity of infection as mentioned in the figures and/or figure legends. PM_10_ treated A549 cells were washed by 1X PBS (phosphate-buffered saline) solution and infected with appropriate RNA viruses in serum-free media as per the subsequent experiment then after 60 minutes, virus containing media was removed from the cells and cells were washed once with 1X PBS solution. Then cells were again supplemented with new PM_10_ containing DMEM media for 24 hours. Samples were then harvested and forwarded for respective quantitative analysis.

### Sampling of airborne particulate matter

Bangalore is an inland city (12°58′ N, 77°34′) situated on the south-central part of India at a height over 900m above sea level. General sources of airborne particulate matter (PM) in the city include vehicular emissions, industrial emissions and re-suspended road dust (http://www.cpcbenvis.nic.in/envis_newsletter/Air%20Quality%20of%20Delhi.pdf; https://www.teriin.org/sites/default/files/2018-08/Report_SA_AQM-Delhi-NCR_0.pdf; http://164.100.107.13/Bangalore.pdf). Air samples were collected from six ambient air quality monitoring sites of Karnataka State Pollution Control Board (KSPCB). Particulate matter with aerodynamic diameter less than 10µm was collected using high volume samplers (Poll tech, India). The samples were collected on quartz fiber filter paper (GE healthcare, India). The filter papers were de-pyrogenated and conditioned prior to sampling [66]. To ensure contamination free sampling, field blanks were included in the samples. After sampling, filter papers were sealed in de-pyrogenated aluminium foil and transported to the laboratory. The samples were stored at - 20°C until further processing. PM on the filter was extracted into methanol. Further, methanol was purged and samples were reconstituted with DMSO [67, 68]. Samples were pooled and used for further experiments.

### Particulate Matter (PM_10_) dose standardization

For all the preliminary experiments three different dosage form of PM_10_ was used in the ratios 1:1 (PM_10_: DMEM), 0.2:1 (PM_10_: DMEM) and 0.5:1 (PM_10_: DMEM) named as PM(I), PM(II) and PM(III) respectively. And after the standardization through different experiments PM(I) that is 1:1 (PM_10_: DMEM) dosage of PM_10_ was used for subsequent experiments.

### SEM-EDS Analysis

Particulate Matter (PM) dissolved in appropriate solvents was installed on the metallic stabs in the form of droplets and dried overnight in the desiccator for complete solvent dry process. Samples were then loaded on the high-resolution field emission scanning electron microscope (SEM) (HR FESEM) from Zeiss, model name ULTRA Plus at IISER Bhopal for PM_10_ morphological analysis. Then chemical composition of the PM_10_ was elucidated by the Energy Dispersive X-ray spectrometer (EDS) component of the scanning electron microscope.

### Quantitative real-time reverse transcription PCR

Total RNA was extracted with the Trizol reagent (Ambion/Invitrogen) and used to synthesize cDNA with the iScript cDNA Synthesis Kit (BioRad, Hercules, CA, USA) according to the manufacturer’s protocol. Gene expression was measured by quantitative real-time PCR using gene-specific primers and SYBR Green (Biorad, Hercules, CA, USA). The 18S gene was used as a reference control. Real time quantification was done using StepOne Plus Real time PCR Systems by Applied BioSystems (Foster City, CA, USA).

### Luciferase Reporter assays

A549 cells (5 × 10^4^) were seeded into a 12-well plate and transiently transfected with 50 ng of the transfection control pRL-TK plasmid (*Renilla* luciferase containing plasmid) and 200 ng of the luciferase reporter plasmid (*Firefly* luciferase containing plasmid) of IFNβ and ISRE promoters. After 12 hours cells were treated with PM_10_ in the ratio 1:1 (PM_10_: DMEM) and Blank as a control for 24 hours. Then after cells were infected with NDV (MOI 2) for 24 hours. The cells were lysed at 24 hours after final infection, and finally the luciferase activity in total cell lysates was measured with Glomax (Promega, Madison, WI, USA).

### Enzyme-linked immunosorbent assay (ELISA)

A549 cells were treated with PM_10_ in the ratio 1:1 (PM_10_: DMEM) and Blank as a control after 24 hours of seeding. The culture media were harvested at 36 hours after particulate matter treatment and were analysed by specific ELISA kits (Becton Dickinson) according to the manufacturer’s instructions to determine the amounts of *IL6* that were secreted by the cells.

### Cell count Trypan Blue assay

A549 cells were seeded and after 24 hours treated with PM_10_ and blank for 24 hours before NDV infection. Cell supernatant were collected after 36 hours of infection, mixed with trypan blue dye (Sigma) in the ratio 1:1. The mixture then used for counting the dead cells under the microscope.

### Microscopy

A549 cells were seeded along with cover slips in low confluency and next day treated with PM_10_ at a dosage of 1:1 [PM: DMEM] for 24 hours prior to virus infection. Cells were then infected with NDV-GFP (3 MOI) in serum free media for 1 hour. After infection cells were again supplemented with complete media and treated with PM_10_ at a dosage of 1:1(PM_10_: DMEM) for 24 hours at 37°C, 5% CO_2_. Cells were then washed twice with PBS for 5 minutes and fixed in 4% PFA for 20 minutes again washed in PBS and incubated with DAPI (20 mg/ml) for 30 minutes at room temperature and finally washed thrice with PBS. Cover slips then containing cells were carefully mounted on to the glass slides using Fluoroshield (Sigma) as mounting media. Slide was then kept for few hours for drying before imaging. Images were visualized at 40X with Apotome – AXIO fluorescence microscope by Zeiss.

### NGS Analysis

Total RNA was extracted using TRIzol reagent (Ambion/Invitrogen) and assessed for quality. The RNA-Seq paired end libraries were prepared from the QC passed RNA samples using Illumina Trueseq stranded mRNA sample prep kit. Libraries were sequenced using NextSeq500 with a read length (2×75bp), by Eurofins Genomic India Private Limited, India. The Raw reads were assessed for quality using FastQC (Andrews S et al, 2010). The filtering of reads and the removal of adapters were performed using the tool Trimmomatic [69]. Approximately 18 million base pair reads were mapped to the human transcriptome (hg38), using Kallisto [70] and the abundance of the assembled coding transcriptome were projected as transcripts per million (TPM). The transcripts level abundance counts were converted into gene-level abundance counts using the R package, Tximport [71]. Differential expression analysis was performed using Limma package [72]. The genes which were differentially expressed (−1.5< Log FC <1.5) were selected and the gene ontology analysis were performed using DAVID tool [73]. Bubble plots, circle plot, chord plots were generated from the gene ontology and pathway enrichment results generated by DAVID tool, using the R package GOplot [74].

### Statistical analysis

All experiments were carried out along with the appropriate controls, indicated as untreated/untransfected cells (Ctrl) or transfected with the transfection reagent alone (Mock). Experiments were performed in duplicates or triplicates for at least two or three times independently. GraphPad Prism 5.0 (GraphPad Software, La Jolla, CA, USA) was used for statistical analysis. The differences between two groups were compared by using an unpaired two-tailed Student’s t-test. While the differences between three groups or more were compared by using analysis of variance (ANOVA) with Tukey test. Differences were considered to be statistically significant when *P* < *0.05*. Statistical significance in the figures is indicated as follows: *\*\*\*P* < *0.001, **P* < *0.01, *P* < *0.05; ns*, not significant.

## Acknowledgments

We greatly acknowledge Dr. Gangamma S. for providing particulate matter (PM_10_) in its isolated form collected from the city of Bengaluru. We thank Director, ICAR-NIHSAD for providing BSL-3 facility to conduct H5N1 experiments. We express our humble gratitude towards Dr. Santhalembi Chingtham – for infecting the cells with Influenza (H5N1) virus in BSL-3 core facility at ICAR - NIHSAD Laboratory. We are grateful to Indian Institute of Science Education and Research (IISER) Bhopal for providing the Central Instrumentation Facility. We thank all the members of the laboratory of immunology and infectious disease biology for helpful discussions. We acknowledge shutterstock.com for an image.

## Data and materials availability

The NGS (RNA-Sequencing) data for expression profiling reported in this paper have been deposited in the GenBank database (accession no. yet to receive from NCBI).

## Supplementary Figure Legends

**Supplementary Figure 1:**
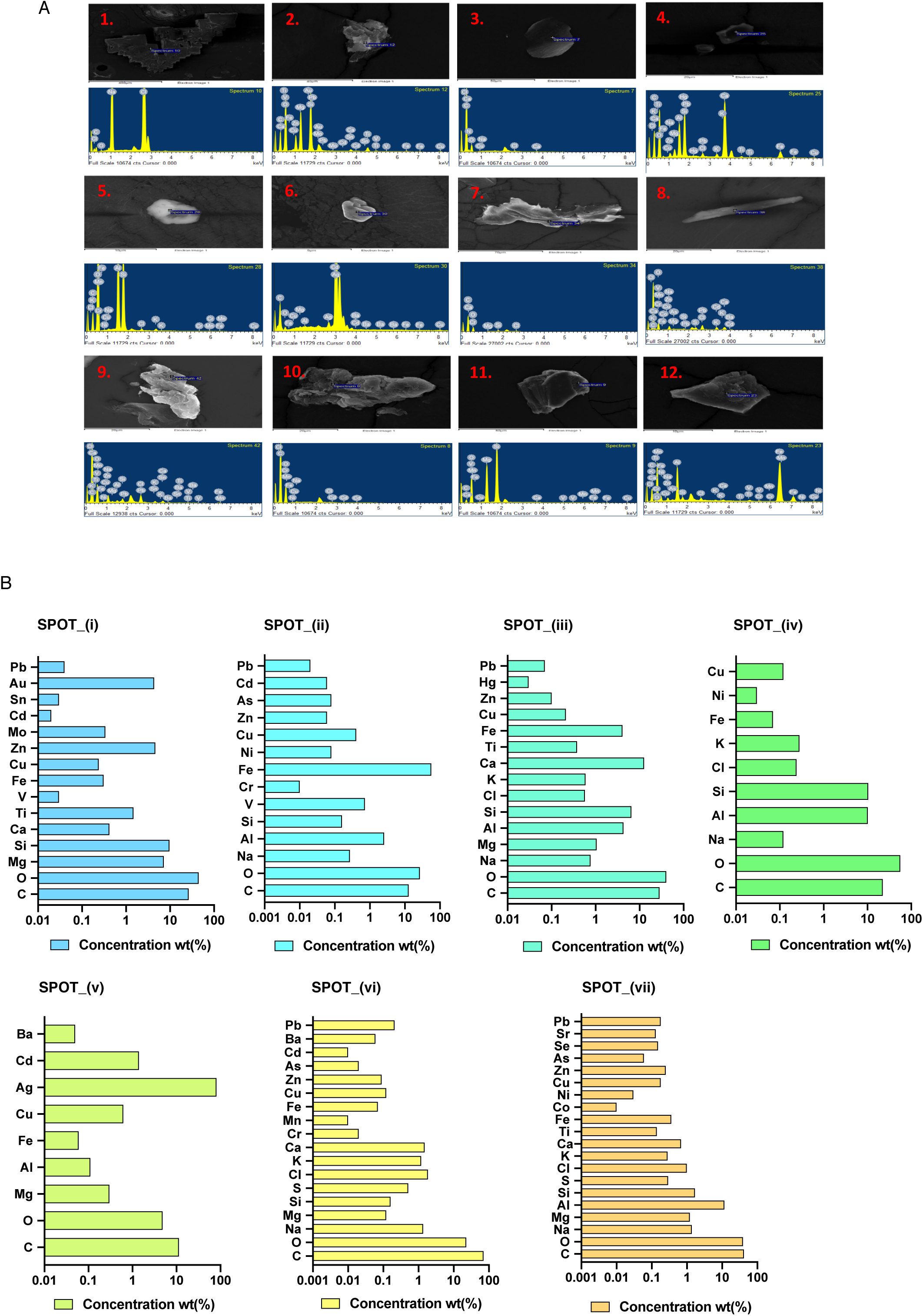
SEM-EDS analysis of PM_10_. Scanning electron images and energy – dispersive X-ray spectra of coarse airborne particulate matter PM_10._ (A) 12 different spots of PM_10_ shows 12 different types of spectral peaks corresponding to presence of specific elements at that point. (B) Representation of elemental composition (% weight) of PM_10_ at few other spots in bar graph having metal name on *y-axis* and respective concentration (%weight) on *x-axis*.

**Supplementary Figure 2:**
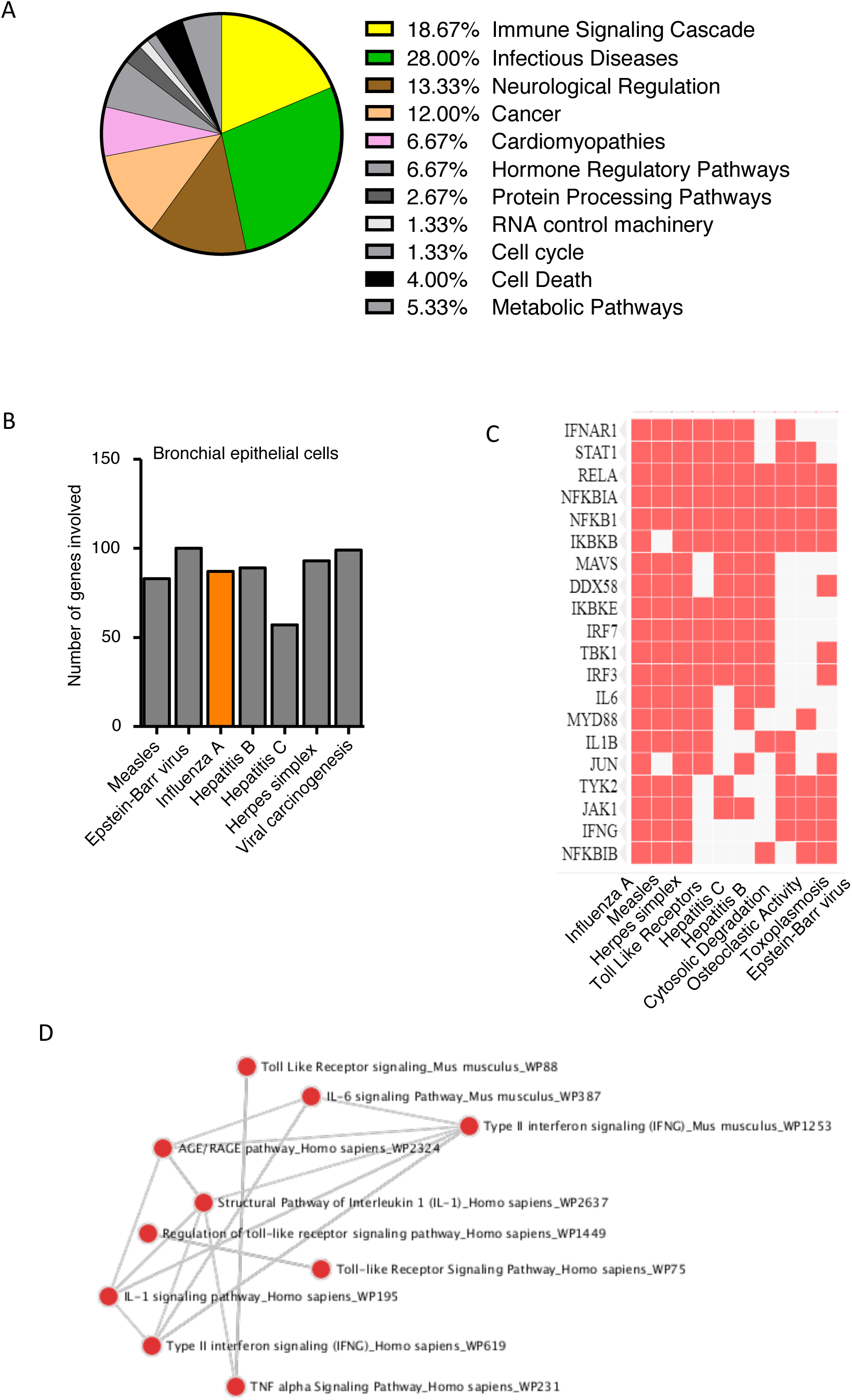
GEO dataset GSE27973 re-analysis. (A) Cellular pathways dysregulated in presence of CSE (cigarette smoke extract) exposure and rhinovirus infection in human bronchial epithelial cells. (B) Number of genes involved in various diseases. (C) Exact gene plotted against the disease in which it is involved, represented in the heat map generated by the Enrichr software. (D) Connecting network between the pathways dysregulated, represented in the dot network analysis generated by the Enrichr software.

**Supplementary Figure 3:**
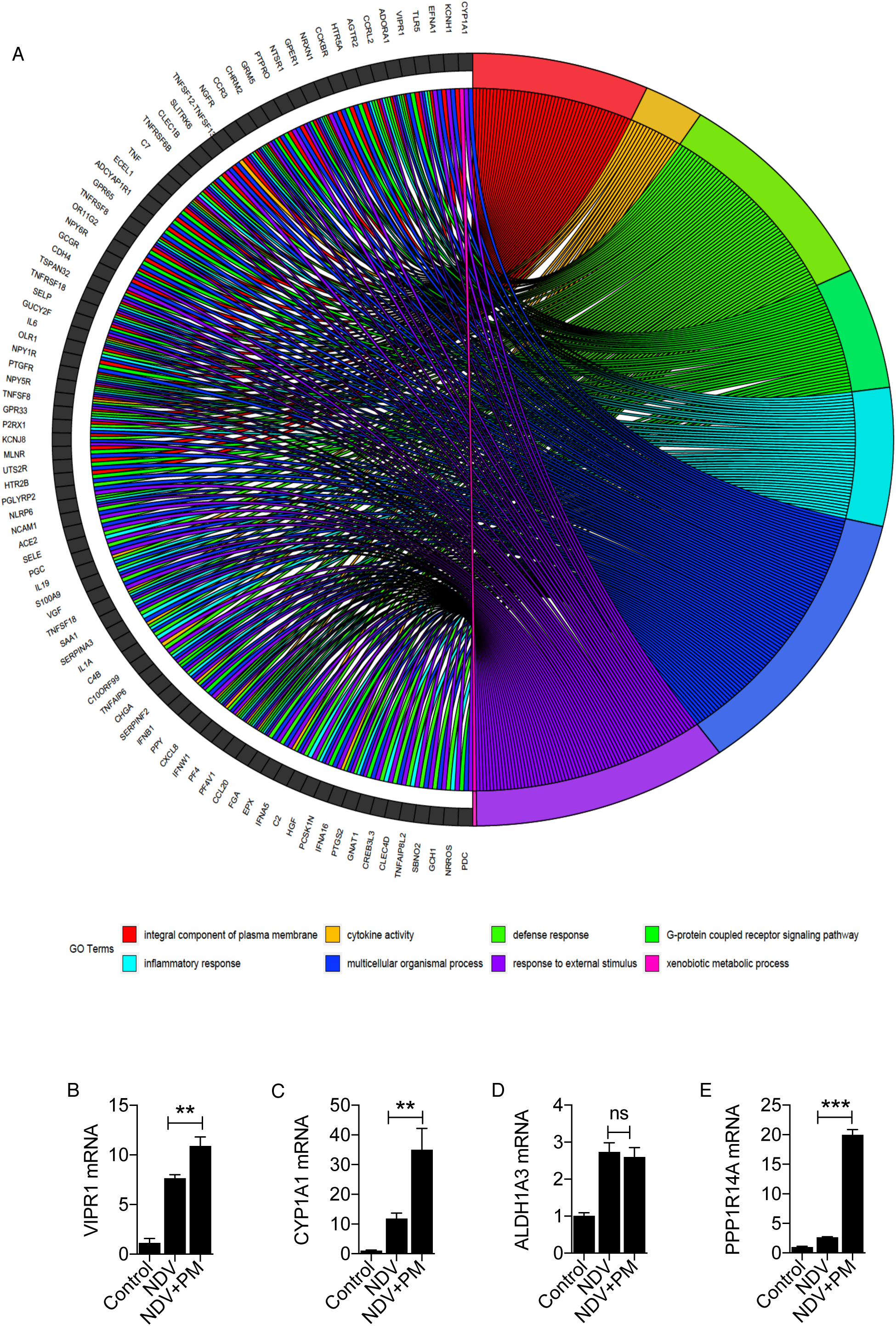
Gene Ontology analysis of PM_10_ treated and H5N1 infected A549 cells. (A) Gene Ontology analysis represented by the chord plot that connects the common differentially expressed genes with the top significantly enriched ontology terms. (B-E) Quantification (measured by qRT-PCR) of the fold changes in the abundances of metabolic pathways related transcripts: VIPR1, CYP1A1, ALDH1A3 and PPP1R14A in A549 cells exposed with PM_10_ and infected with NDV. Sample labelled as untreated (control), mock NDV infected (NDV) and PM_10_ exposed plus NDV infected (NDV+PM). Data are mean +/- SEM of triplicate samples from single experiment and are representative of two independent experiments. \*\*\**P*<0.001, \*\**P*<0.01 and ns = non-significant by one-way ANOVA Tukey test and unpaired t-test.

**Supplementary Figure 4:**
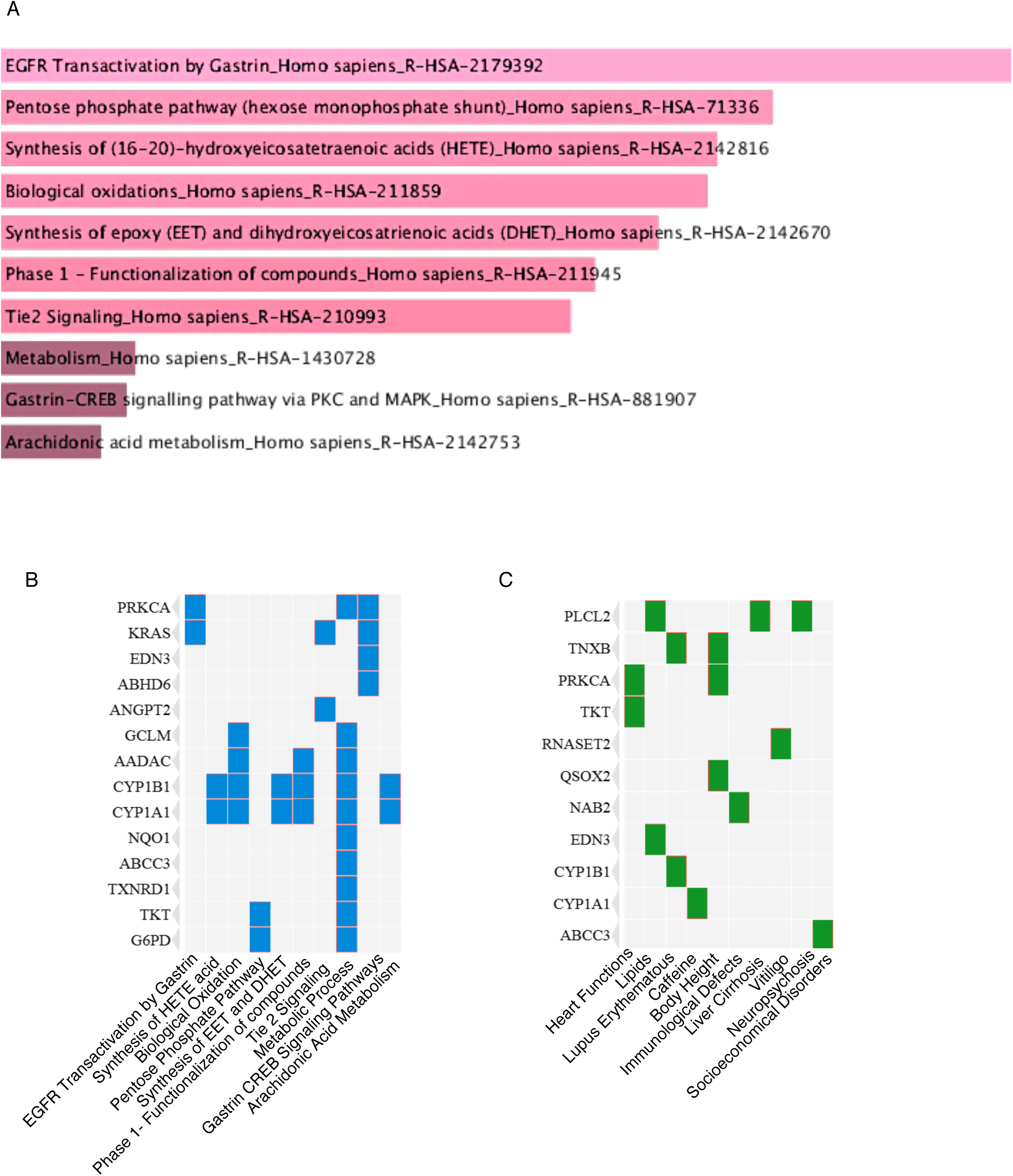
GEO dataset GSE27973 re-analysis to depict the pathways and genes upregulated in presence of CSE and RV infection. Enrichr software is used for the depiction of related genes and pathways. (A) Representation of enriched pathways by bar graph. (B) Representation of upregulated genes involved in enriched pathways by heat map. (C) Representation of upregulated genes involved in various diseases by heat map. Here, human bronchial epithelial cell lines were exposed to CSE (cigarette smoke extract), RV (rhinovirus).

